# Do *Brucella* Antibody Titers Correlate with Clinical Outcomes and Culture Positivity of Brucellosis?

**DOI:** 10.1101/534941

**Authors:** Shahad A. Alsubaie, Shouq A. Turkistani, Alanoud A. Zeaiter, Abrar K. Thabit

**Affiliations:** Faculty of Pharmacy, King Abdulaziz University, Jeddah, Saudi Arabia

**Keywords:** *Brucella*, brucellosis, serology, antibody, titer, culture, Saudi Arabia

## Abstract

Brucellosis is a zoonotic disease caused by *Brucella* spp., namely *B. meletinsis* and *B. abortus* in humans. Studies on the correlation between *Brucella* antibody titers and clinical outcomes are limited. Therefore, this study assessed such correlation and evaluated the correlation between baseline serologic results with culture positivity and clinical picture. Patients tested positive for *Brucella* antibodies at baseline and diagnosed with brucellosis between January 2008 and December 2018 were included. Collected data included clinical outcomes, baseline culture positivity, arthralgia, baseline and EOT temperature, white blood cell (WBC) count, C-reactive protein level, and erythrocyte sedimentation rate. Of 695 patients tested for *Brucella* antibodies, only 94 had positive baseline serology and diagnosed with brucellosis, among whom 63 had EOT serology. No significant correlations were found between EOT antibody titers of both *Brucella* spp. and clinical cure, mortality, length of stay, and duration of therapy. Additionally, no correlations were found between baseline serology and culture positivity, arthralgia, temperature, and other lab values. *Brucella* serology does not correlate with clinical outcomes at EOT nor with culture positivity at baseline. Therefore, healthcare providers are advised to consider the whole clinical picture of a brucellosis patient without relying solely on serologic results during follow up and not replace culturing with serology testing alone at the time of diagnosis.

## INTRODUCTION

*Brucella* spp. is a Gram-negative coccobacillus that causes Brucellosis (also known as “Malta fever”); a contagious zoonotic infectious disease (1). The genus *Brucella* consists of seven species, four of which causes a disease in humans. The most common species include *B. melitensis* which is transmitted from goat and sheep followed by *B. abortus* that comes from cows. *B. suis* has a more restricted occurrence and is transmitted from pigs. Lastly, *B. canis* is uncommonly associated with human diseases but can be transmitted from dogs (2, 3). Brucellosis patients usually acquire the disease through consumption of unpasteurized dairy products or direct contact with infected animals or their fluids (4). The infection usually starts with nonspecific symptoms, such as fatigue, headache, dry cough, and night sweats (2). However, in severe cases it can be associated with variable complications (2).

To aid the clinical diagnosis of brucellosis, laboratory methods are utilized which include direct and indirect methods. Direct methods include microbiological culturing as well as the detection *Brucella* gene using polymerase chain reaction (PCR) test (1). Brucella culture is considered the gold standard and provides the definitive diagnosis of the disease (5). However, as *Brucella* is a slow-growing bacterium, it may take the culture a week or longer to show growth (6). Additionally, patients with a long-standing disease may have negative culture results. Therefore, indirect serological assays are used alternatively and provide faster results. The Rose Bengal testing, tube agglutination, and enzyme-linked immunosorbent assay (ELISA) are the most recommended methods of *Brucella* serology (1). Moreover, some nonspecific laboratory tests, such as white blood cell (WBC) count, C-reactive protein (CRP), erythrocyte sedimentation rate (ESR), serum lactate dehydrogenase, and alkaline phosphatase may show some changes (6).

In a response to an infection with *Brucella* spp., the immune system naturally produces antibodies. IgM isotype is first produced followed by the IgG and IgA isotypes. With appropriate treatment, the titer of these antibodies should gradually fall off. The remaining high titer of IgG and IgA indicate a possibility of relapse or progress to a chronic focal disease (2).

A retrospective study of brucellosis patients showed that *Brucella* antibodies remained persistently positive even after the patients demonstrated full clinical recovery and cure (7). As serological cure of brucellosis is defined when antibody titers reach less than 1:320, the rate of serological cure in this study increased from 8.3% in the first three months to 71.4% after two or more years with a median time for a serological cure of 18.5 ± 3 months. In addition, none of the 116 patients included in the study experienced a relapse at the end of the follow up period although some patients did not achieve the serological cure by this time point. Besides this study, no other studies addressing the relationship between *Brucella* antibody titers and clinical outcomes were found in the literature. As such, this study aims to evaluate such correlation, as well as to assess the correlation of baseline serology with baseline culture positivity and clinical picture in an area of disease endemicity.

## METHODS

### Study design and patients

This was a retrospective, cross-sectional study of brucellosis patients admitted to King Abdulaziz University Hospital in Jeddah, Saudi Arabia during the period from January 2008 to December 2018. Patients aged 18 years or older who had positive *Brucella* serology (regardless of the titer), diagnosed with brucellosis based on serology (defined as antibody titer of ≥ 1:640), *Brucella* culture, or clinical picture, and received antibiotic treatment were included in the study. Patients who did not receive treatment for brucellosis despite positive serology (of < 1:640) were excluded as they may not have received the diagnosis of the disease.

Given the endemic nature of brucellosis in Saudi Arabia, an antibody titer of at least 1:640 of either *Brucella* spp. is needed to confirm the serological diagnosis, especially in the absence of positive *Brucella* culture or classic clinical symptoms suggestive of brucellosis.

### Endpoints

The primary endpoint was the correlation of *Brucella* antibody titers with clinical cure at the end of therapy (EOT). Secondary endpoints included the correlations of EOT *Brucella* antibody titers with mortality, length of stay, duration of therapy, and EOT temperature, WBC count, CRP level, and ESR. Additional secondary endpoints were the correlations of baseline *Brucella* serological results with *Brucella* culture positivity, baseline temperature, WBC count, CRP level, and ESR.

### Statistical analysis

Descriptive statistics using percentages, mean ± standard deviation for normally distributed data, and median (interquartile range, IQR) for non-normally distributed data were used. Normal distribution was determined using Shapiro-Wilk test for normality. Pearson’s correlation was used to assess the correlation between *Brucella* antibody titers with different outcomes at EOT and baseline. Two-tailed statistical significance was indicated by a *P* value of < 0.05. SPSS version 24.0 (SPSS, Inc., Chicago, Illinois, USA) was used for statistical analysis.

## RESULTS

Out of 695 patients who tested for *Brucella* via serology or culture, only 94 had positive serology and were diagnosed with brucellosis. Table 1 lists the characteristics of patients included in the study. More than half of the patients were males with an average age of 48 ± 19 years. Overall, patients did not have fever or leukocytosis at baseline; though, CRP levels and ESR were mostly elevated. In addition, less than one-third of the patients had complicated brucellosis or arthralgia. Sixty three of the 94 patients included in the study had EOT serological results available. Median antibody titers at EOT were lower than their baseline counterparts. Similarly, median CRP level and ESR showed a decrease from baseline. The majority of patients experienced clinical cure and only 5 died.

**Table 1.** Patients characteristics

| Characteristic | All Patients<br>(n=94) |
| --- | --- |
| Age, years (mean ± SD) | 48 ± 19 |
| Sex, male, n (%) | 58 (61.7) |
| Race, n (%)<br>• White<br>• Black<br>• Asian | 83 (88.3)<br>3 (3.2)<br>8 (8.5) |
| Location, n (%)<br>• Outpatient<br>• Inpatient medical ward<br>• ICU | 43 (45.7)<br>49 (52.1)<br>2 (2.1) |
| Charlson co-morbidity index (mean ± SD) | 1.9 ± 2.1 |
| Baseline *B. melitensis* antibody titer (mean IQR) | > 1:1280 1:1280->1:1280 |
| Baseline *B. melitensis* antibody titer ≥ 1:640, n (%) | 78 (83) |
| Baseline *B. abortus* antibody titer (median IQR) | > 1:1280 1:1280->1:1280 |
| Baseline *B. abortus* antibody titer ≥ 1:640, n (%) | 73 (77.7) |
| Baseline temperature, °C (median IQR) | 36.8 36-37.3 |
| Baseline white blood cells count, cells/mm<sup>3</sup> (mean ± SD) | 6.6 ± 4.9 |
| Baseline C-reactive protein, mg/L (mean ± SD) | 34.5 ± 39.9 |
| Baseline erythrocyte sedimentation rate, mm/hr (mean ± SD) | 29 ± 23.5 |
| Duration of therapy (median IQR) | 45 45-135 |
| Risk factors for infection, n (%) |  |
| • Unpasteurized dairy product<br>• Direct contact with animals<br>• Both<br>• Other<br>• Unknown | 42 (44.7)<br>4 (4.3)<br>8 (8.5)<br>1 (1)<br>39 (41.5) |
| Diagnostic test positivity, n (%) |  |
| • *Brucella* serology alone<br>• Both *Brucella* serology and culture | 42 (44.7)<br>52 (55.3) |
| *Brucella* spp., n (%) |  |
| • *B. melitensis*<br>• *B. abortus*<br>• Both | 4 (4.3)<br>2 (2.1)<br>88 (93.6) |
| Complicated brucellosis, n (%) | 29 (30.9) |
| • Spondylodiscitis<br>• Neurobrucellosis<br>• Orchitis<br>• Other<br>• > 1 organ | 17 (59)<br>5 (17)<br>3 (10.3)<br>3 (10.3)<br>1 (3.4) |
| Arthralgia, n (%) | 30 (31.9) |
| Regimen used, n (%) |  |
| • Aminoglycoside + doxycycline + rifampin<br>• Doxycycline + rifampin<br>• Other | 44 (46.8)<br>34 (36.2)<br>16 (17) |
| Length of stay, days (median IQR)<sup>a</sup> | 9 5-17 |
| Clinical cure, n (%) | 80 (85.1) |
| Mortality, n (%) | 5 (5.3) |
| EOT *B. melitensis* antibody titer (median IQR)<sup>b</sup> | 1:320 1:320-1:640 |
| EOT *B. melitensis* antibody titer < 1:320, n (%)<sup>b</sup> | 28 (29.8) |
| EOT *B. abortus* antibody titer (median IQR)<sup>b</sup> | 1:320 1:160-1:640 |
| EOT *B. abortus* antibody titer < 1:320, n (%)<sup>b</sup> | 33 (35.1) |
| EOT temperature, °C (median IQR) | 36.7 36.5-36.9 |
| EOT white blood cells count, cells/mm<sup>3</sup> (mean ± SD) | 6.2 ± 3.4 |
| EOT C-reactive protein, mg/L (median IQR) | 3.2 3.1-3.3 |
| EOT erythrocyte sedimentation rate, mm/hr (median IQR) | 8 5-22 |

EOT, end of therapy; IQR, interquartile range
<sup>a</sup> Data for hospitalized patients only.
<sup>b</sup> Only 63 of patients had end of therapy serology and hence the percentage reflects this population.

The correlations between EOT *Brucella* antibody titers with different outcome variables are shown in Table 2. Table 3 shows the correlation after stratifying the antibody titers based on the serological cure criterion of < 1:320. A significant, though weak, correlation between clinical cure and low EOT antibody titers of both *Brucella* spp. was found. Nevertheless, this correlation was lost when antibody titers were stratified based on the cutoff value of 1:320. This can be further recognized in Figure 1 where almost equal rates of clinical cure and failure was observed in the subgroup of patients who had EOT antibody titers of < 1:320. Furthermore, no significant correlations were found between EOT antibody titers of either *Brucella* spp. and other outcomes except for EOT WBC count where lower *B. abortus* antibody titers only were weakly associated with higher WBC counts (negative correlation). Interestingly, such correlation turned towards the opposite direction after the stratification based on the serological cure cutoff. In addition of both of these correlations with EOT WBC counts being weak as represented by the low correlation coefficient, patients generally had normal mean WBC count at this time point. Subgroup analyses of patients with uncomplicated or complicated brucellosis showed similar results of lack of correlation between the EOT antibody titers and all of the tested variables.

**Figure 1.**
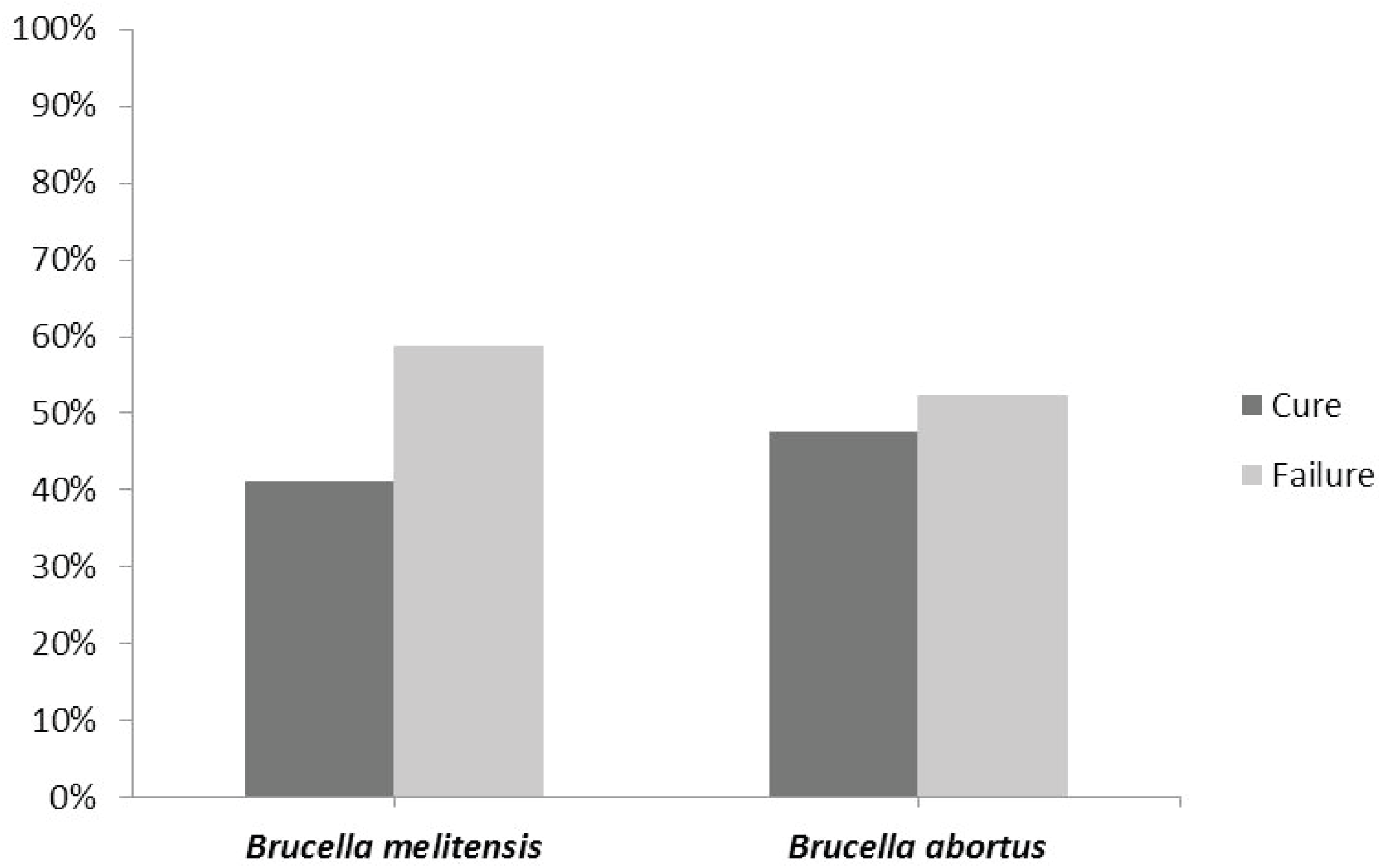
Distribution of clinical outcomes in patients with end of therapy antibody titers of < 1:320

**Table 2.**
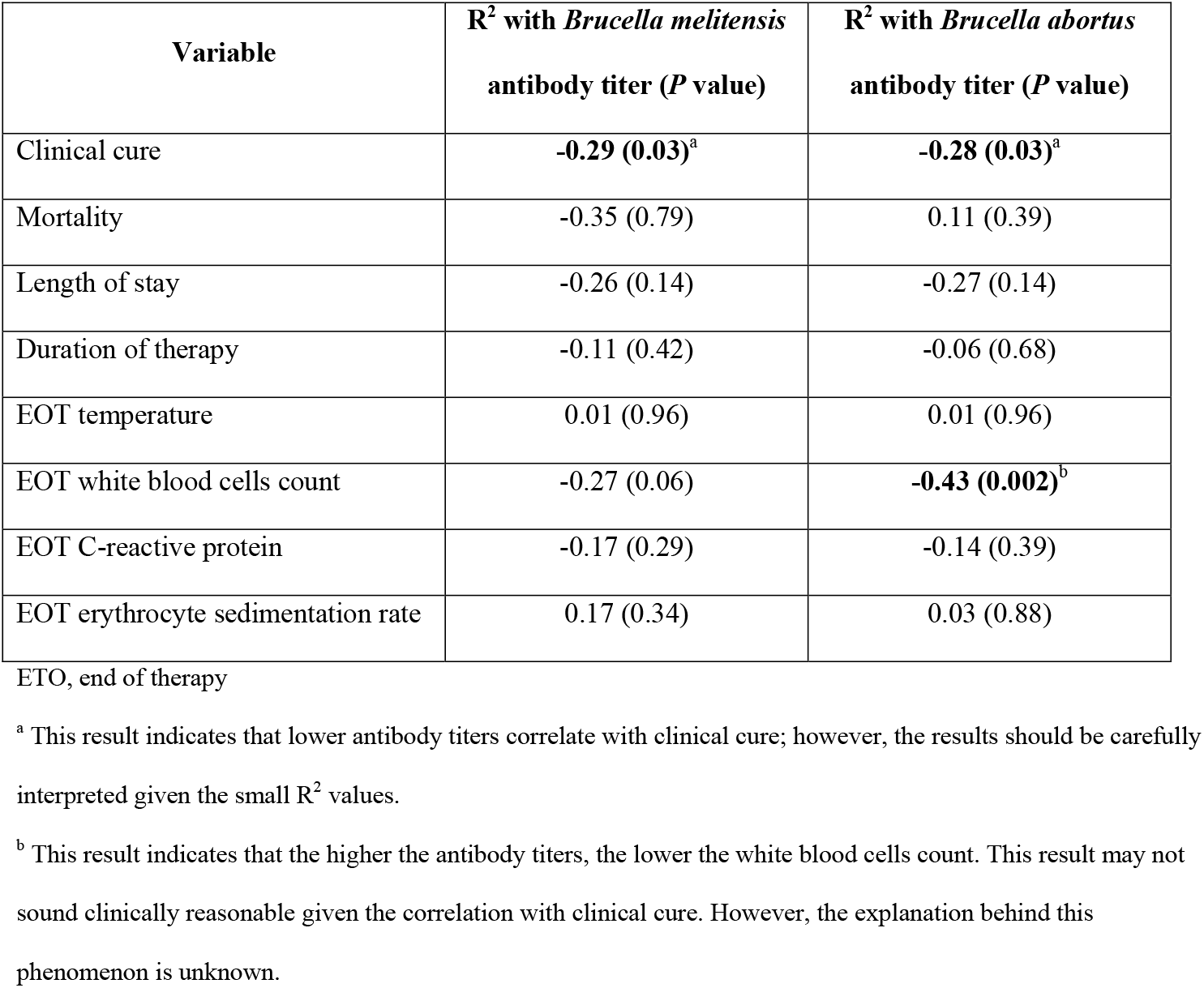
Pearson’s correlation coefficient of the correlations between end of therapy *Brucella* antibody titers with different outcome variables (n=63)

| Variable | R<sup>2</sup> with *Brucella melitensis*<br>antibody titer (*P* value) | R<sup>2</sup> with *Brucella abortus*<br>antibody titer (*P* value) |
| --- | --- | --- |
| Clinical cure | **-0.29 (0.03)<sup>a</sup>** | **-0.28 (0.03)<sup>a</sup>** |
| Mortality | -0.35 (0.79) | 0.11 (0.39) |
| Length of stay | -0.26 (0.14) | -0.27 (0.14) |
| Duration of therapy | -0.11 (0.42) | -0.06 (0.68) |
| EOT temperature | 0.01 (0.96) | 0.01 (0.96) |
| EOT white blood cells count | -0.27 (0.06) | **-0.43 (0.002)<sup>b</sup>** |
| EOT C-reactive protein | -0.17 (0.29) | -0.14 (0.39) |
| EOT erythrocyte sedimentation rate | 0.17 (0.34) | 0.03 (0.88) |

EOT, end of therapy
<sup>a</sup> This result indicates that lower antibody titers correlate with clinical cure; however, the results should be carefully interpreted given the small R<sup>2</sup> values.
<sup>b</sup> This result indicates that the higher the antibody titers, the lower the white blood cells count. This result may not sound clinically reasonable given the correlation with clinical cure. However, the explanation behind this phenomenon is unknown.

**Table 3.** Pearson’s correlation coefficient of the correlations between end of therapy *Brucella* antibody titers of < 1:320 with different outcome variables (n=63)

| Variable | R<sup>2</sup> with *Brucella melitensis*<br>antibody titer < 1:320 (*P*<br>value) | R<sup>2</sup> with *Brucella abortus*<br>antibody titer < 1:320 (*P*<br>value) |
| --- | --- | --- |
| Clinical cure | 0.19 (0.14) | 0.16 (0.2) |
| Mortality | 0.02 (0.89) | -0.01 (0.93) |
| Length of stay | 0.25 (0.16) | 0.35 (0.05) |
| Duration of therapy | 0.12 (0.39) | 0.09 (0.53) |
| EOT temperature | -0.04 (0.83) | 0.09 (0.62) |
| EOT white blood cells count | 0.09 (0.52) | **0.33 (0.02)<sup>a</sup>** |
| EOT C-reactive protein | 0.26 (0.12) | 0.19 (0.24) |
| EOT erythrocyte sedimentation rate | -0.29 (0.1) | -0.16 (0.38) |

EOT, end of therapy
<sup>a</sup> This result indicates that antibody titers of ≥ 1:320 are correlated with lower white blood cells count. As indicated in Table 2, this result may not sound clinically reasonable given the correlation with clinical cure. However, the explanation behind this phenomenon is unknown.

Table 4 and Table 5 list the correlation results of different baseline variables with baseline antibody titers before and after stratification based on the serological criterion for diagnosis of ≥ 1:640, respectively. While it seems that the antibody titers of *B. abortus* correlate with culture positivity in both tables, this correlation can be considered weak with the low value of the correlation coefficient, as well as by visualizing the lack of difference in the rate of culture positivity among patients who met the serological diagnostic criterion at baseline (Figure 2).

**Figure 2.**
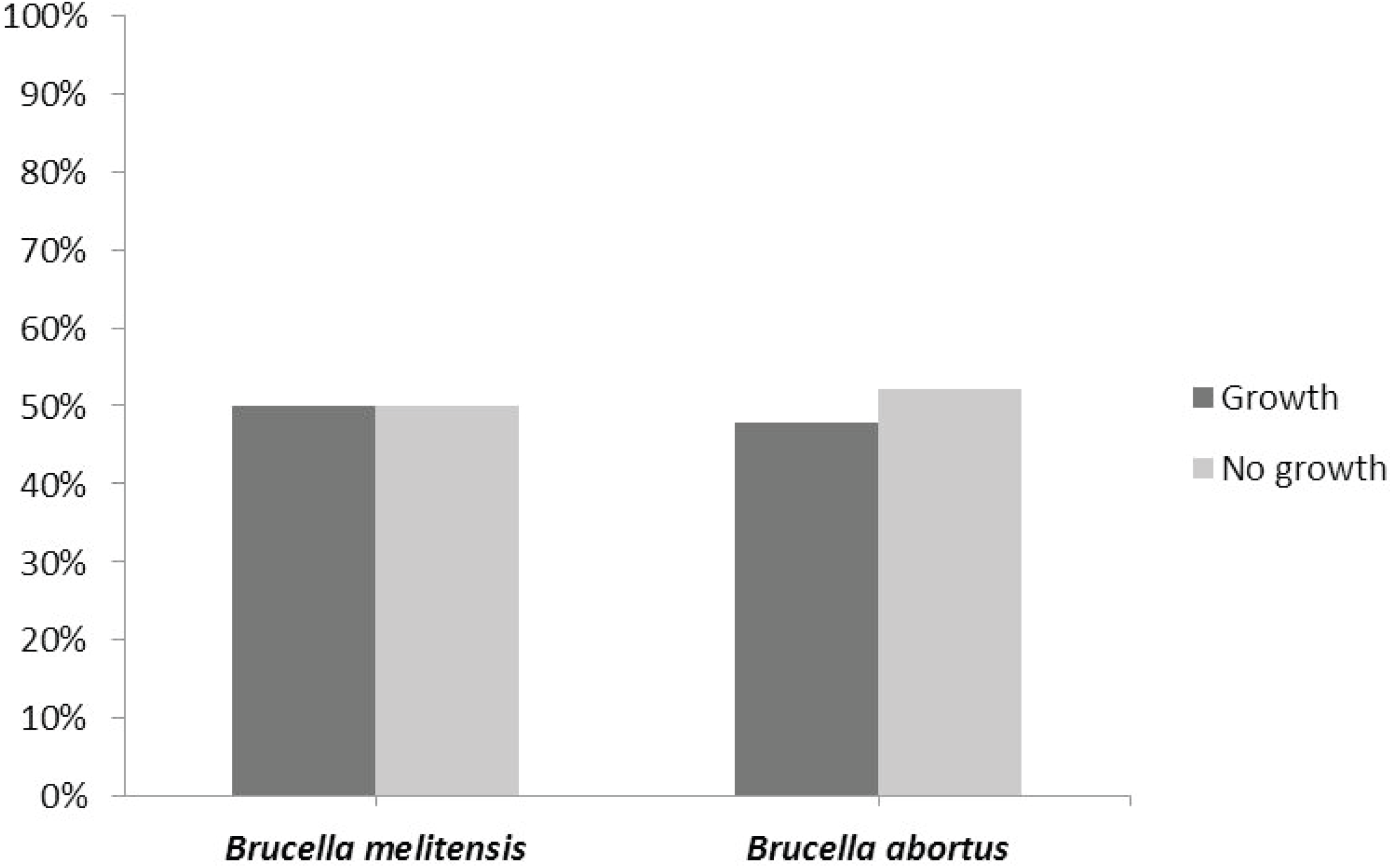
Distribution of culture results in patients with baseline antibody titers of ≥ 1:640

**Table 4.** Pearson’s correlation coefficient of the correlations between baseline *Brucella* antibody titers with different baseline variables (n=94)

| Variable | R<sup>2</sup> with *Brucella melitensis*<br>antibody titer (*P* value) | R<sup>2</sup> with *Brucella abortus*<br>antibody titer (*P* value) |
| --- | --- | --- |
| Age | -0.05 (0.65) | -0.04 (0.71) |
| Location | 0.04 (0.73) | 0.2 (0.05) |
| Culture positivity | 0.14 (0.17) | **0.29 (0.004)<sup>a</sup>** |
| Arthralgia | 0.02 (0.86) | 0.09 (0.4) |
| Complicated brucellosis | -0.09 (0.37) | **-0.25 (0.02)<sup>b</sup>** |
| Baseline temperature | 0.07 (0.84) | -0.002 (0.99) |
| Baseline white blood cells count | 0.03 (0.77) | -0.01 (0.92) |
| Baseline C-reactive protein | 0.03 (0.79) | 0.04 (0.74) |
| Baseline erythrocyte sedimentation<br>rate | -0.001 (0.99) | -0.01 (0.97) |

<sup>a</sup> This result indicates that the higher the antibody titers, the higher the probability that a culture is positive in the 56.4% of the total patient population who had positive *Brucella* culture at baseline.
<sup>b</sup> This result indicates that the complicated brucellosis was associated with lower antibody titers at baseline. This result should be carefully interpreted given the low correlation coefficient.

**Table 5.** Pearson’s correlation coefficient of the correlations between baseline *Brucella* antibody titers of ≥ 1:640 with different baseline variables (n=94)

| Variable | R<sup>2</sup> with *Brucella melitensis*<br>antibody titer $\geq 1:640$ (*P*<br>value) | R<sup>2</sup> with *Brucella abortus*<br>antibody titer $\geq 1:640$ (*P*<br>value) |
| --- | --- | --- |
| Age | -0.03 (0.78) | -0.11 (0.31) |
| Location | -0.01 (0.9) | 0.17 (0.1) |
| Culture positivity | 0.19 (0.7) | **0.3 (0.002)<sup>a</sup>** |
| Arthralgia | 0.02 (0.86) | 0.1 (0.33) |
| Complicated brucellosis | 0.03 (0.78) | -0.15 (0.14) |
| Baseline temperature | 0.21 (0.87) | 0.03 (0.82) |
| Baseline white blood cells count | -0.03 (0.75) | -0.08 (0.46) |
| Baseline C-reactive protein | 0.02 (0.9) | 0.03 (0.8) |
| Baseline erythrocyte sedimentation<br>rate | 0.06 (0.7) | -0.06 (0.69) |

<sup>a</sup> This result indicates that the higher the antibody titers, the higher the probability that a culture is positive in the 56.4% of the total patient population who had positive *Brucella* culture at baseline.

In addition to the lack of correlation between baseline antibody titers of either species with age and location, no correlations were found with the clinical manifestations of the disease. While complicated brucellosis seems to be related with a lower antibody titer at baseline as given by the significant *P* value, the correlation was considered null as it became insignificant when the antibody titers were divided into two groups based on the cutoff value of ≥ 1:640. In both cases, it should be noted that the correlation coefficient was low.

## DISCUSSION

Brucellosis is an endemic disease common in Saudi Arabia, Iraq, Jordan, and other Middle East countries, as well as Turkey, central Asia, Mexico, South America, and Asia-Pacific region (8, 9). As brucellosis is endemic, only few studies exist in the literature addressing the condition, and studies concerning antibodies and their relationship with disease outcomes are even scarce. Hence, this study aimed to fill the gap by assessing this relationship.

In a study of 116 clinically cured brucellosis patients, Almuneef, et al found that *Brucella* antibodies remained detectable at end of therapy follow up despite full clinical recovery (7). However, at a later time point after two years post completion of therapy, the antibody titers decreased dramatically and reached undetectable levels in some patients. A similar finding was seen in our patient population at EOT where about half of the patients who experienced clinical success did not achieve the serological success of antibody titers of < 1:320. This fact is also supported by the lack of correlation between the antibody titers and clinical outcomes, including temperature and infection markers (WBC count, CRP level, and ESR). Nevertheless, it is presumed that *Brucella* antibody titers of those patients would eventually reach undetectable levels after 24 months or more after the EOT.

Our study also looked at the correlation of *Brucella* antibody titers with the clinical picture of brucellosis including the most common symptom of arthralgia, as well as fever, leukocytosis, and elevated inflammatory markers, CRP and ESR. A lack of correlation was demonstrated between *Brucella* antibody titers and these variables. These results resemble the findings from a study by Alsubaie and colleagues who evaluated the family members of brucellosis patients whether they appeared symptomatic or asymptomatic (10). Symptoms sought in this study included arthralgia, fever, malaise, headache, anorexia, and weight loss. WBC counts and inflammatory markers were not assessed. Forty of the 178 family members screened for brucellosis had classic manifestations of the disease including arthralgia and fever, of whom only 18 (45%) had positive serology results (≥ 1:160). Thus, they were confirmed to have acute brucellosis (five had positive serology tests but claimed to have past infections). Such result further confirms the lack of correlation between brucellosis symptoms and serology test for *Brucella* as the remaining 55% symptomatic members had negative serology tests. This lack of correlation was also observed when only four (3%) of 138 members had positive serology tests despite absence of symptoms, hence they were diagnosed with the disease (seven had past infections with positive serology results).

To our knowledge, this is the first study to evaluate the correlation between antibody titers and culture positivity. Since our institution is located in an endemic country, patients are serologically diagnosed with brucellosis when antibody titers measure 1:640 or higher. However, when this criterion is not matched, a patient may still be diagnosed with the disease based on a positive blood culture for *Brucella* or classic clinical picture of the disease. As demonstrated in Figure 2, a positive serology does not always come with a positive culture and vice versa. In the study by Alsubaie, et al, only 8 (47.1%) of 17 seropositive patients had positive cultures for *B. melitensis* vs. none in the asymptomatic brucellosis group (10). Both the findings of this study and the findings of our study indicate that serology testing for *Brucella* should not replace culturing, the gold standard testing for the disease. Furthermore, when serological results at baseline were assessed against the factors of patient’s age and hospital location in our study, a correlation was not observed.

As studies on the relationship of serological test results of brucellosis with clinical outcomes and culture positivity are limited, this study comes to provide an evidence that *Brucella* serology does not correlate with clinical outcomes at EOT nor with culture positivity at baseline. Therefore, healthcare providers are advised to consider the complete clinical picture of a brucellosis patient, as well as culture, without relying solely on serologic results.

## Acknowledgements

We would like to thank Rawan O. Al-Madfaa, Mai A. Alalawi, Lana O. Basudan, Shahad F. Alhejaili for their help with data collection. Also, we would like to thank the IT department and the immunology lab of King Abdulaziz University Hospital for providing the lists of patients.

